# A Real Time Quaking Induced Conversion (RT-QuIC) Assay for Detection of Misfolded Insulin Protein

**DOI:** 10.64898/2026.04.08.717350

**Authors:** Jessica M. Alderiso, Rafael Hernandez LaTorre, Trisha M. Cox, Matthew G. DiGiovanni, Keileigh Fulbright, Brenda F. Canine

**Affiliations:** Touro University College of Osteopathic Medicine-Montana; Weissman Hood Institute at Touro University, McLaughlin Research Institute

**Keywords:** insulin, misfolded protein, aggregation, protein folding, RT-QuIC

## Abstract

Protein misfolding plays a critical role in aging and disease, yet the involvement of specific proteins in metabolic dysfunction is still poorly understood. Here, we report studies on the development of a Real-time Quaking-Induced Conversion (RT-QuIC) assay to detect misfolded insulin, a peptide hormone required for blood glucose regulation. Although RT-QuIC assays were originally designed to amplify misfolded prion proteins implicated in neurodegeneration, we adapted the method to monitor conformational changes in insulin. We first validated the RT-QuIC insulin assay using recombinant insulin and insulin aggregates recovered from clinical infusion devices. Protein characterization by gel electrophoresis, circular dichroism, and particle size analysis suggests differences in insulin recovered from the infusion device. We then applied the RT-QuIC assay to tissue samples from a mouse model of metabolic disease. This work provides proof-of-concept of a novel assay for studying the role of insulin aggregation in disease progression and aging. The RT-QuIC assay for insulin may also provide new avenues to explore early detection, mechanistic insights, and therapeutic targets of metabolic disorders linked to aging and disease.

## 1. Introduction

Metabolic dysfunction is a driver of many age-related diseases including Type 2 Diabetes Mellitus (T2DM) [1–4]. Regulation of glucose levels under homeostasis has been described in detail [5]; essentially, increased glucose levels in the blood trigger the release of insulin from the beta cells in the pancreas which triggers release of insulin into the blood stream where it interacts with receptors on tissues. Insulin interaction with its receptors consequently begins a signaling cascade which opens a glucose transporter to enable entry of glucose into the cells of the tissues and the glucose can then be used in many processes including cellular respiration which results in ATP production [5]. In T2DM, the pathway of insulin release and glucose uptake is disrupted leading ultimately to a feedback loop where more glucose stays in the blood, so more insulin is released, and so forth, leading to hyperinsulinemia and the state of insulin insensitivity or insulin resistance [3,6]. Risk factors that contribute to insulin resistance in addition to aging include obesity, inactivity, and liver dysfunction [7]. However, the underlying mechanisms that cause or contribute to insulin resistance are poorly understood. Moreover, therapeutics such as metformin and semaglutide do not work on insulin resistance directly, but instead metformin works on the liver and inhibits gluconeogenesis, whereas semaglutide stimulates insulin secretion [8,9]. Side effects of these drugs are also very significant [10]. Of the over $49 trillion dollars spent on healthcare in the US (17.6% of total GDP), one in four dollars is estimated to be spent on care for diabetics [11]. Understanding the underlying root mechanisms of insulin resistance and dysfunction is a critical undertaking if we are to develop more effective, targeted, and/or patient-specific therapeutics and early diagnostics.

Real-time quaking-induced conversion (RT-QuIC) assays are a powerful protein amplification and detection technique developed originally to quantify the formation of misfolded prion aggregates [12–15]. RT-QuIC leverages the ability of a misfolded insulin protein seed to convert recombinant insulin protein monomers into a β-sheet-rich structure, a process monitored by the binding of a fluorescent dye, such as thioflavin T (ThT). Researchers then monitor and record the protein misfolding process as outlined in numerous studies [14–30]. Over the years, technological advancements, such as optimization of substrates and reagents, have significantly improved the sensitivity and reproducibility of RT-QuIC assays [15,27].

RT-QuIC assays have been successfully applied to animal and human neurodegenerative conditions, including sporadic and inherited Creutzfeldt-Jakob disease, fatal familial insomnia, Gerstmann-Sträussler-Scheinker syndrome, Parkinson’s disease, Alzheimer’s disease and amyotrophic lateral sclerosis (ALS) [16,18–22,27,30] (**Table 1**). Before the development of RT-QuIC assays, most studies of neurodegenerative diseases relied almost entirely on post-mortem brain tissue to identify hallmark protein aggregation in histological pathologies. More recently, increasing evidence points to a broader role for protein misfolding and aggregation in systemic diseases [31–33]. Particularly, T2DM has been suggested to involve structural abnormalities of endogenous or recombinant insulin [33]. Nonetheless, traditional methods often fail to detect insulin protein misfolding effectively due to their poor sensitivity, limited reproducibility, and lack of scalability.

**Table 1.** Summary of Prion and Protein dysfunction diseases.

| **Table 1. Summary of Prion and Protein dysfunction diseases.** |  |  |  |
| --- | --- | --- | --- |
| **Misfolded Protein** | **Disease Model(s) Studied** | **Sample Type(s)** | **Study** |
| **Prion protein (PrP)** | Fatal Familial Insomnia (FFI)<br>Gerstmann-Straußler-Scheinker (GSS)<br>Creutzfeldt Jakob Disease (CJD) | CSF | Rhoads et al., 2020 |
| **PrP** | CJD | CSF, nasal | Orrú et.al. 2015 |
| **PrP** | CJD | CSF, olfactory mucosa | Bongianni et. al. 2017 |
| **PrP** | CJD | CSF | Lattanzio et. al. 2017 |
| **PrP** | CWD | Saliva | Henderson, et. al 2013 |
| **Alpha-synuclein (α-Syn)** | Parkinson's Disease (PD), Dementia with Lewy Bodies (DLB), Alzheimer's Disease (AD), Progressive Supranuclear palsy (PSP) | Brain, CSF | Manne et. al. 2019 |
| **PrP** | CJD, Scrapie, CWD, Bovine Spongiform Encephalopathy (BSE), GSS, FFI | Brain, CSF | Atarshi et. al 2011 |
| **PrP, α-Syn, tau** | CJD, Pick Disease, PD, DLB | Brain, CSF | Saijo et. al 2018 |
| **α-Syn** | PD, DLB, Multiple Systems Atrophy, Pure Autonomic Failure |  | Rossi e. al. 2020 |
| **Super Oxide Dismutase I (SOD-1)** | Familial amyotrophic Lateral Sclerosis (fALS), sporadic amyotrophic Lateral Sclerosis (sALS) | Brain | Mielke et. al. 2024 |
| **PrP** | sCJD | CSF | Naranjo et. al. 2025 |
| **PrP** | CWD | Soil | Grunklee et. al. 2025 |
| **PrP** | CWD | Deer lymph nodes | Darish et. al. 2025 |
| **α-Syn** | DLB, PD | Brain, intestinal mucosa, skin | Parveen, et.al. 2025 |

As a review, insulin has 2-subunits and 51 amino acids and is critical for maintaining glucose homeostasis in the body (**Figure 1**). Insulin signaling is a tightly regulated process that enables the body to maintain glucose homeostasis, primarily by promoting glucose uptake in muscle and adipose tissue and suppressing glucose production in the liver. Briefly, as diagramed in **Figure 1D**, insulin is a peptide hormone secreted by pancreatic beta cells in response to elevated blood glucose levels [34]. Insulin binds to the insulin receptor (IR) on the surface of target cells, including muscle, liver, and adipose cells. Insulin binds to the alpha subunit of the IR, causing a conformational change, activating the tyrosine kinase activity of the receptor. The activated receptor phosphorylates the IR sub-strate proteins, insulin receptor substrate 1 (IRS1) and insulin receptor substrate 2 (ISR2), which, once phosphorylated, serve as docking sites for downstream molecules, namely phosphoinositide 3-kinase (PI3K). PI3K converts phosphatidylinositol 4,5, bisphosphate 2 (PIP2) to phosphatidylinositol 3,4,5-triphosphate (PIP3) at the inner plasma membrane, resulting in the recruitment of 3-phophoinosiditide-dependent kinase -1 (PDK1) and protein kinase B (Akt). Akt then promotes the translocation of GLUT4 (glucose transporter 4) to the cell membrane, allowing for cellular glucose uptake. Akt also inhibits gluconeogenesis and promotes glycogen synthesis in the liver while also stimulating lipogenesis and inhibiting lipolysis in adipose tissue. Insulin activates the mechanistic target of rapamycin (mTOR) pathway, which regulates cell proliferation and gene expression (**Table 2**).

**Table 2.**
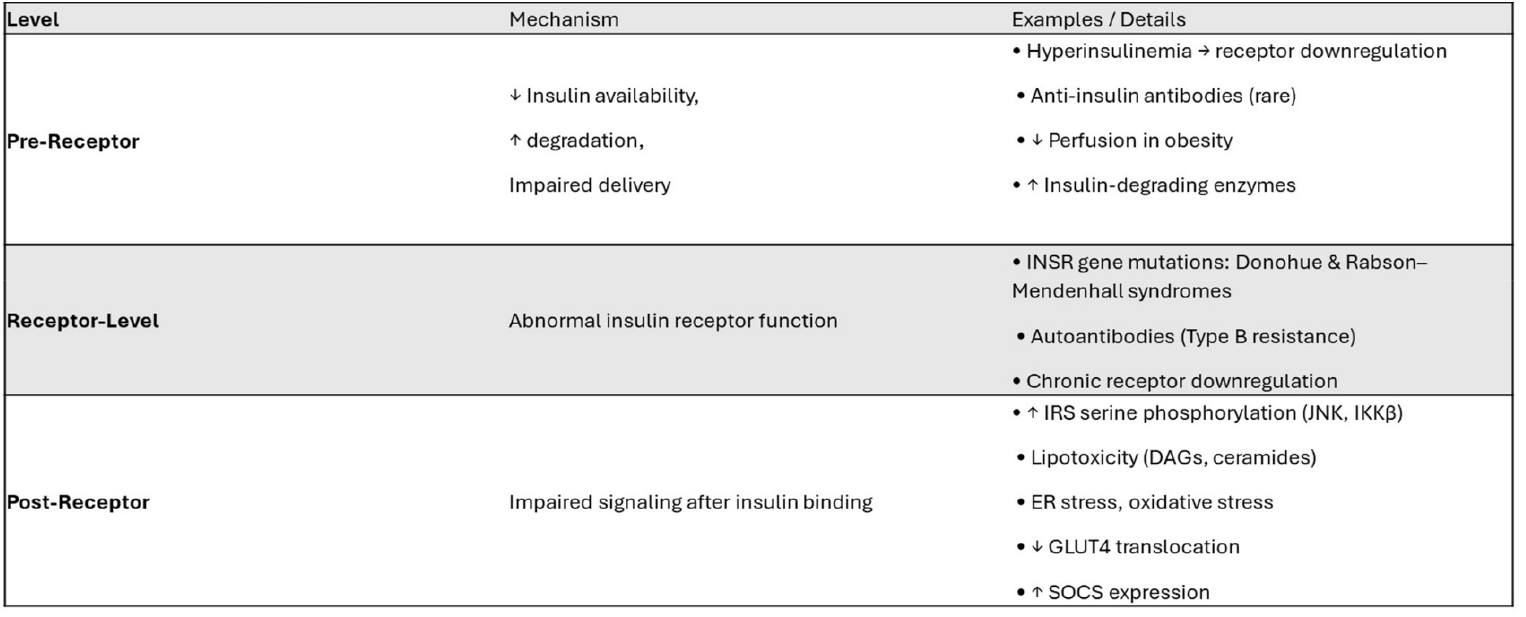
Summary of proposed mechanisms for insulin resistance.

**Figure 1:**
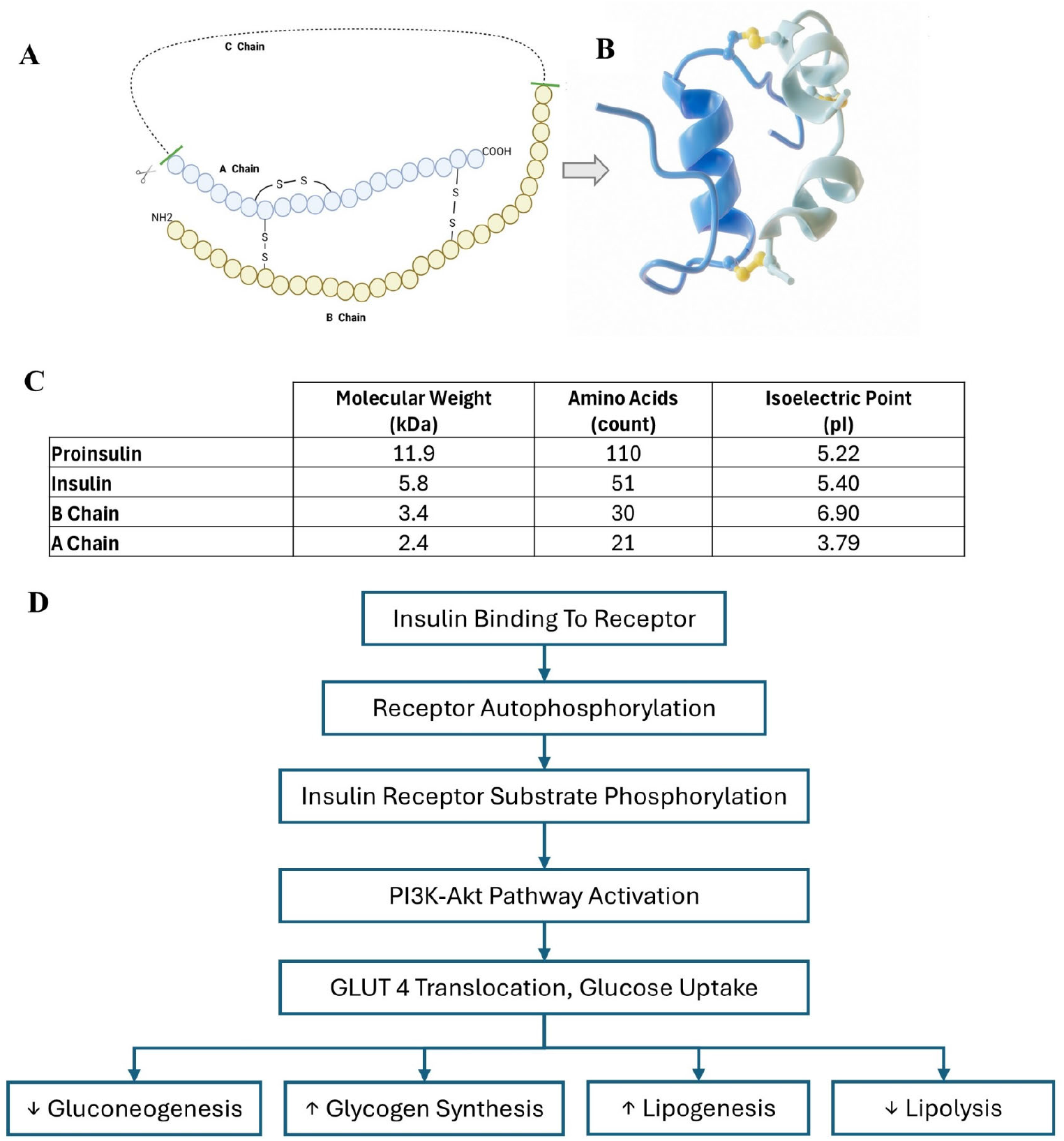
Structural and biochemical properties of proinsulin, insulin and subunits. **A)** Amino acid diagram showing the key activating step in insulin hormone function involves cleavage of proinsulin, resulting in the release of C-peptide. a hallmark of activated endogenous insulin in circulation. **B)** A ribbon diagram of the functionally active mature form of insulin consisting of an alpha and beta chain. **C)** Properties of combined and independent components of the insulin protein. The molecular weight and number of amino acids is reduced during the proinsulin cleavage event to insulin, with a corresponding change in isoelectric point (pI). **D)** The signaling cascade of insulin interacting with the membrane insulin receptor is diagrammed and ultimately leads to translocation of GLUT4 and increases in glucose uptake. This shifts the metabolism to glycogenesis (glycogen synthesis) and lipogenesis and inhibits gluconeogenesis and lipolysis.

Insulin-derived amyloidosis is characterized by structural, kinetic, and thermodynamic alterations leading to insulin aggregation and is a recognized complication associated with insulin injections, significantly impacting insulin absorption and action [35,36]. Insulin-derived amyloidosis is sometimes described as an ʺinsulin ball” and may be characterized by increasing insulin resistance due to the presence of insulin-positive amyloid masses often at the injection sites [37]. It has also been suggested that insulin and proinsulin misfolding may be an early event in the progression to T2DM and a target for intervention [38–41]. It is also well established that, according to the manufacturer’s storage instructions, recombinant exogenously delivered insulin should be stored in a mesothermic temperature range, avoiding both cold and hot conditions [42]. Experts recommend discarding any insulin in plastic pump cartridges or tubing after 48 hours due to a decrease in efficacy. Insulin resistance can arise from various mechanisms, such as at the pre-receptor, receptor, and post-receptor levels, including impaired insulin availability, receptor downregulation, and altered intracellular signaling pathways (**Table 2**). Based on these observations, we hypothesized that RT-QuIC could be adapted to detect misfolded insulin aggregates in metabolic dysfunction. Here, we present data on the development of an RT-QuIC seed amplification assay to measure insulin seeding activity. The assay uses recombinant insulin substrate and detects the ability of misfolded insulin seeds to induce conformational conversion of the normally folded recombinant insulin. This conversion is monitored as a structural change from the native alpha-helical form to a beta-sheet rich amyloid conformation detectable by fluorescence of ThT providing a readout of insulin amyloid-forming activity.

## 2. Materials and Methods

### Protein Preparation and Insulin Aggregates

We used commercially available recombinant human insulin (Humulin R, Eli Lilly) as the substrate for the RT-QuIC assay. To collect aggregated insulin, we recovered insulin from occluded insulin pump cartridges and tubing (T: slim X2 3 mL Cartridge, Tandem Diabetes Care) and used the aggregated insulin as seed for the RT-QuIC assay. The recovered aggregated insulin seeds were aliquoted into 1 μL volumes (equivalent to 0.1 standard insulin units) and stored at 4°C until use. Protein concentrations were estimated using standard insulin unit conversion metrics and determined by spectrophotometric analysis (A280) using a NanoDrop™ spectrophotometer (ThermoFisher) with the protein specific extinction coefficient applied.

### SDS-PAGE and Silver Staining

We prepared non-aggregated and aggregated insulin samples in Tricine SDS sample buffer (Thermo Fisher Scientific), with Sample Reducing agent (Thermo Fisher Scientific). Reduced samples were heated at 95°C for 5 minutes, and then briefly centrifuged. We loaded 10 μg of each sample onto a 10–20% Tricine SDS-PAGE gel (NuPAGE, Thermo Fisher Scientific), alongside a Spectra Multicolor Low Range Protein Ladder (Thermo Fisher Scientific), and ran the gel at 120 V for 60 minutes in 1X Tricine SDS running buffer. Following electrophoresis, the Tricine-SDS-PAGE gel was stained and developed using a Silver Staining Kit (Thermo Fisher Scientific), according to the manufacturer’s protocol. Images were obtained using a smartphone camera against a white background (Android and Apple).

### Circular Dichroism Spectroscopy

We diluted recombinant insulin in 1X Dulbecco’s Phosphate Buffered Saline (DPBS) pH 7.4 (Gibco) to a concentration of 0.1 mg/mL and acquired circular dichroism spectra in the far-UV (200-260 nm) range using a Jasco J815 spectropolarimeter equipped with a temperature controller and a 1 cm pathlength quartz cuvette. A multi-scan modality was utilized, and an average wavelength scan was collected every 1 nm from 260-200 nm. This method allowed us to visualize secondary structural change shifts associated with mis-folding. Beta Structure Selection methods (BeStSel), a freely accessible webserver for secondary structure and fold recognition for circular dichroism (CD) spectra was utilized to estimate and characterize the secondary structure. This algorithm was developed specifically to characterize amyloid fibrils by CD spectroscopy. Raw CD data was input into the tool and analyzed for 8 domains of structure. Percent of structure was estimated and then alpha helical structure domains, beta sheet domains, and other (turns, disordered, etc.) are reported.

### Dynamic Light Scattering (DLS)

We analyzed the particle size distribution using a Zetasizer (Malvern Instruments), equipped with a 1 cm quartz flow cell. We diluted recombinant insulin to a final concentration of 3 μg/mL in DPBS (pH 7.4), used a syringe to carefully load 1 mL of the diluted insulin into the flow cell to minimize air bubbles, and recorded three technical replicates per sample. The Zetasizer software (v7.12) calculated the Z-average hydrodynamic diameter and polydispersity index. Statistically significant comparisons were made using a student’s t-test (alpha = 0.005).

### RT-QuIC Reaction Setup

We performed RT-QuIC assays in black 96-well plates with flat clear optical bottoms (Nunc, Thermo Fisher Scientific). Each 150 μL reaction contained 10 mM DPBS (pH 7.4), 10 μM ThT (Sigma-Aldrich) and the protein to be tested for aggregation: either recombinant insulin (2.65 standard units) or an equivalent amount (90 μg) of purified protein namely, Bovine Serum Albumin (Bio-Rad) or Lysozyme (Enzo Life Sciences). Each protein tested for aggregation was also seeded with aggregates: either aggregated recombinant insulin recovered from an insulin pump (0.008 standard units), an equivalent amount of bovine serum albumin or lysozyme (0.29 μg) or tissue homogenate (1 μL) whose preparation is described in the mouse tissue samples methods section.

A master mix of each experimental condition was created then aliquoted into 4 technical replicates in the 96-well plate. Plates were sealed with adhesive film (3M) and incubated at 37°C in a FLUOstar Omega plate reader (BMG Labtech), programmed for double orbital cyclical shaking at 500 RPM for 1-minute, followed by a 1-minute resting period that repeated for the duration of the time course. ThT fluorescence was measured every 15-20 minutes for 50–72 hours using 450 nm excitation and 480 nm emission filters with a gain setting of 900.

### RT-QuIC Data Collection and Analysis

Fluorescence intensity data was analyzed using OMEGA-MARS v5.01 (BMG Labtech) and Microsoft Excel. Reactions were considered positive if ≥ 3 of 4 technical replicates showed a fluorescence intensity that crossed a predefined threshold calculated as the mean fluorescence signal of the negative controls plus five standard deviations (a criterion previously established in prion RT-QuIC methodologies to minimize false-positive detection while maintaining assay sensitivity). Fluorescence curves were inspected to confirm consistent kinetics and to exclude sporadic signal spikes or artifacts. ThT fluorescence (RFUs) mean and standard deviation of 4 technical replicates were calculated and plotted using Microsoft Excel. To quantify the kinetics of the RT-QuIC reactions, half-time was calculated as the time (hours) an experimental condition reached half of its maximum fluorescence; The time which an experimental sample fluorescence increased over the negative controls was subtracted from the time the sample reached is maximum fluorescence and began to plateau, then divided in half. Statistically significant comparisons were made using a student’s t-test (alpha = 0.005).

### Mouse Tissue Samples

Adipose tissues were collected postmortem from J:ARC(S) (034608) mice housed in the McLaughlin Research Institute AALAAC accredited animal facility. The tissues were immediately snap frozen then stored at -80°C until processing. Tissues were weighed then suspended in 1X PBS, pH 7.4 so that the tissue mass was 10% of the total volume. Tissues were homogenized manually using a Dounce homogenizer. Homogenized tissue was then diluted 1:100,000, 1:1,000,000, and 1:10,000,000 in 1X PBS, pH 7.4 then utilized as seed in the previously described RT-QuIC protocol.

## 3. Results

Our goal was to develop and optimize an RT-QuIC assay using recombinant human insulin as a substrate and aggregated insulin as the seed. To validate this assay, we used *in vitro* insulin aggregates as seed material and extended the application to *in vitro* purified proteins and tissue homogenates from a mouse model of neurodegeneration and glucose dysregulation.

### 3.1. Circular Dichroism Shift Signifies Insulin Conformation Shift

Using circular dichroism, we measured ellipticity in the far UV range (200-260 nm) **(Figure 2)**. Aggregated insulin recovered from an insulin pump tubing shows a shift from the non-aggregated recombinant human insulin. A distinct negative signal was observed at 220 nm in the aggregated insulin when compared to a negative peak at 210 nm for non-aggregated insulin. CD spectra analysis using BeStSel modeling estimated a secondary structure composition for non-aggregated insulin of 40.4% α-helix, 0% β-sheet and 59.6% other conformations. Aggregated insulin was modeled to be 26.7% α-helix, 32.1% β-sheet and 41.2% other conformations.

**Figure 2:**
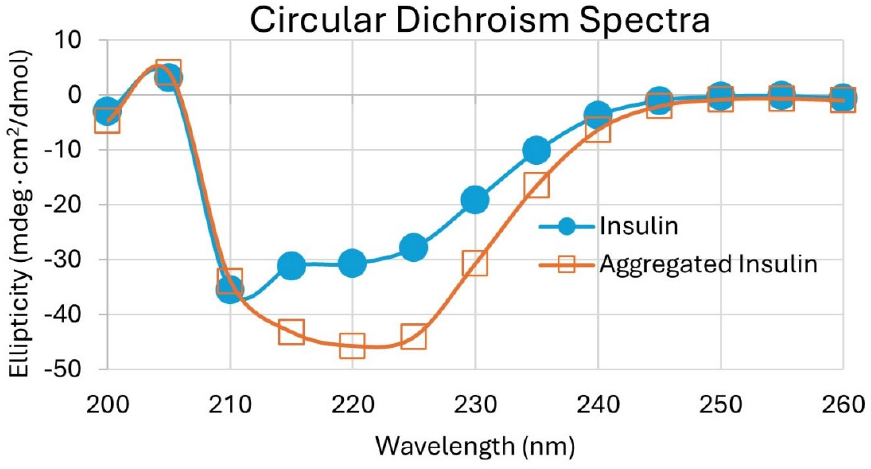
Circular dichroism (CD) spectra of insulin and aggregated insulin protein. Alpha helices show a characteristic negative signal al 210 nm whereas antiparallel beta sheets have a negative peak at 220 nm. CD spectra in the far UV range show a shift from non-aggregated insulin compared to aggregated insulin.

### 3.2 Gel Electrophoresis and Particle Size Analysis Confirms Insulin Aggregation

Characterization of the insulin substrate and seed materials used for the RT-QuIC assay was performed using protein gel electrophoresis and particle sizing. We used protein gel electrophoresis in reducing sample conditions visualized by silver staining to verify the insulin substrate and seed materials (**Supplemental Figure 1**). With reducing agent, 2 protein bands < 5 kDa in size were visible in both non-aggregated and aggregated insulin, indicating the alpha and beta chains of insulin **(Supplemental Figure 1)**.

Particle size analysis (**Figure 3**) was performed using a Zetasizer on non-aggregated insulin from fresh vials and aggregated insulin from a clogged insulin pump. Although most of the aggregated insulin displayed the same diameter and volume distribution as the non-aggregated insulin, a small portion of the aggregated sample showed increased particle diameter and volume, confirming the presence of larger aggregates (p < 0.001).

**Figure 3:**
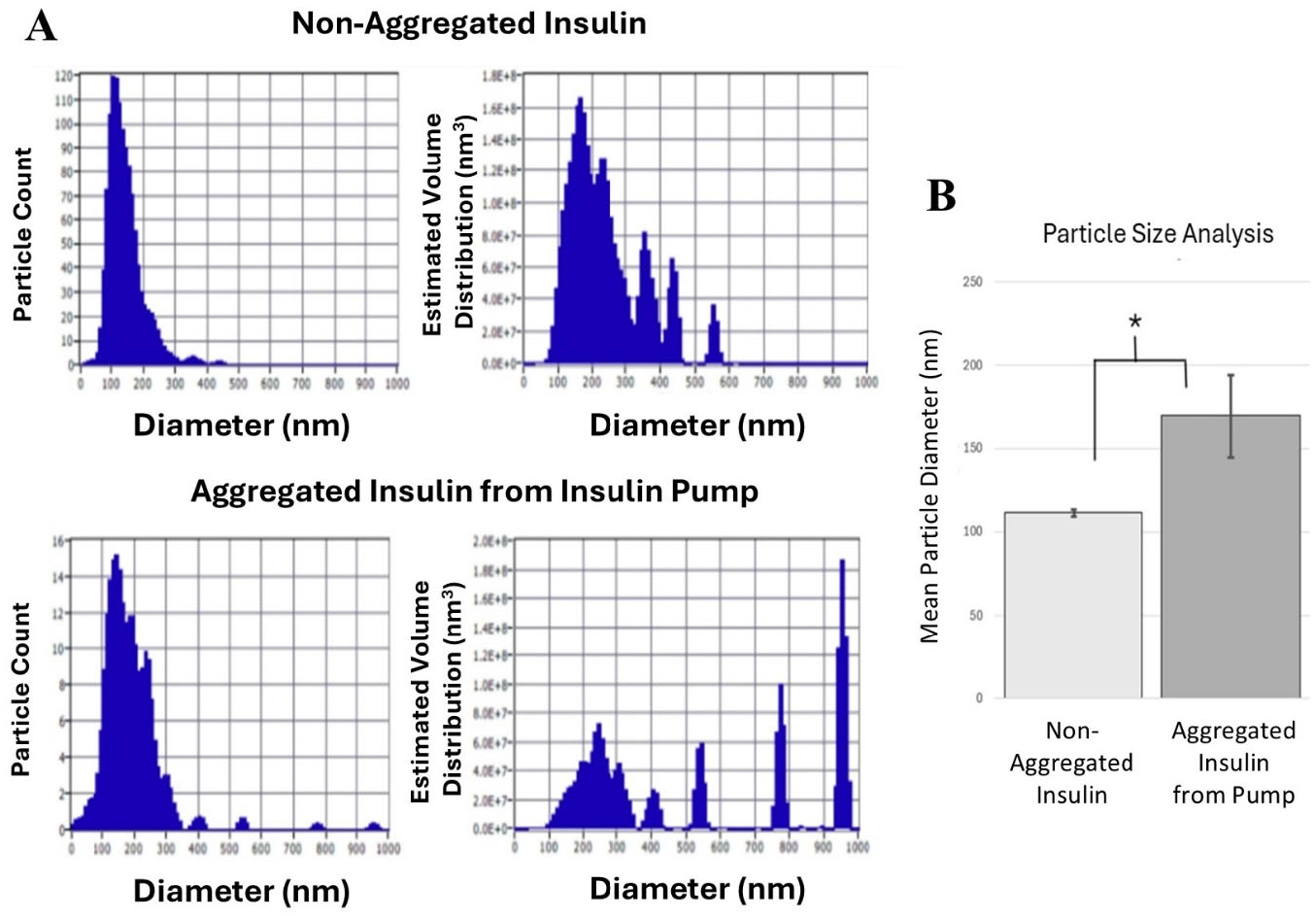
Characterization of insulin and aggregated insulin. **A)** Particle size characterization of insulin and aggregated insulin via use of dynamic light scattering (DLS). Upper panels shows distribution of insulin obtained from a vial (non-aggregated) and bottom panel shows distributions for insulin obtained from insulin pump cartridge (aggregated) Protein samples are diluted in 1X PBS buffer pH 7.4 and loaded into cuvette via syringe through Luer lock port. The particles are visualized on screen and concentration is adjusted to ensure detection of individual particles. Upon measurement a multipoint detection system is used to sample multiple points in the cuvette to for sample distribution. Software then calculates particle number versus diameter (left panels) and particle volume vs diameter (right panel). **B)** Comparison of mean particle diameter ± standard deviation for recombinant insulin obtained from vial (non-aggregated) or from insulin pump cartridge (aggregated). Significant differences were observed in mean size distribution of insulin relative to aggregated insulin using particle size distribution (*p< 0.001).

### 3.3 RT-QuIC Fluorescence Reveals Insulin Seeding Activity In Vitro

We conducted RT-QuIC studies using ThT fluorescence as a reporter for aggregate formation. ThT is weakly fluorescent in solution, but upon binding to beta-rich structures it becomes highly fluorescent. Detection of aggregate formation of commercially available human insulin using RT-QuIC follows the workflow in **Figure 4A. Figure 4B-C** displays single sample curves for visual clarity. Specifically, **Figure 4B** shows RT-QuIC negative controls of buffer only, buffer with fluorescence reporter (ThT), and buffer with ThT fluorescent reporter and the seed. No increase in ThT fluorescence is detected in the negative controls. Experimental RT-QuIC samples comparing 1) non-aggregated insulin with ThT to 2) non-aggregated insulin seeded with aggregated insulin from clogged pump tubing and ThT are shown in **Figure 4C**. Notably, the non-aggregated insulin when subject to the RT-QuIC assay took ∼20 hrs before it began to form insulin aggregates. When aggregated insulin from the clogged cartridge of the infusion pump was included as seed in the RT-QuIC assay, the non-aggregated insulin aggregated much more quickly; in less than 12 hours. Mean ThT fluorescence and standard deviation between replicate samples was calculated and shown in **Supplemental Figure 2A** while **Supplemental Figure 2B** displays individual sample curves. We quantified this difference in kinetics by calculating the time to half-maximum fluorescence (half-time) of insulin aggregation with and without seed aggregates. Samples consisting of insulin with seed aggregates showed significantly faster aggregation (11.3 hour half-time) compared to insulin alone (33.5 hour half-time) (**Supplemental Figure 3, p =0.000189)**.

**Figure 4:**
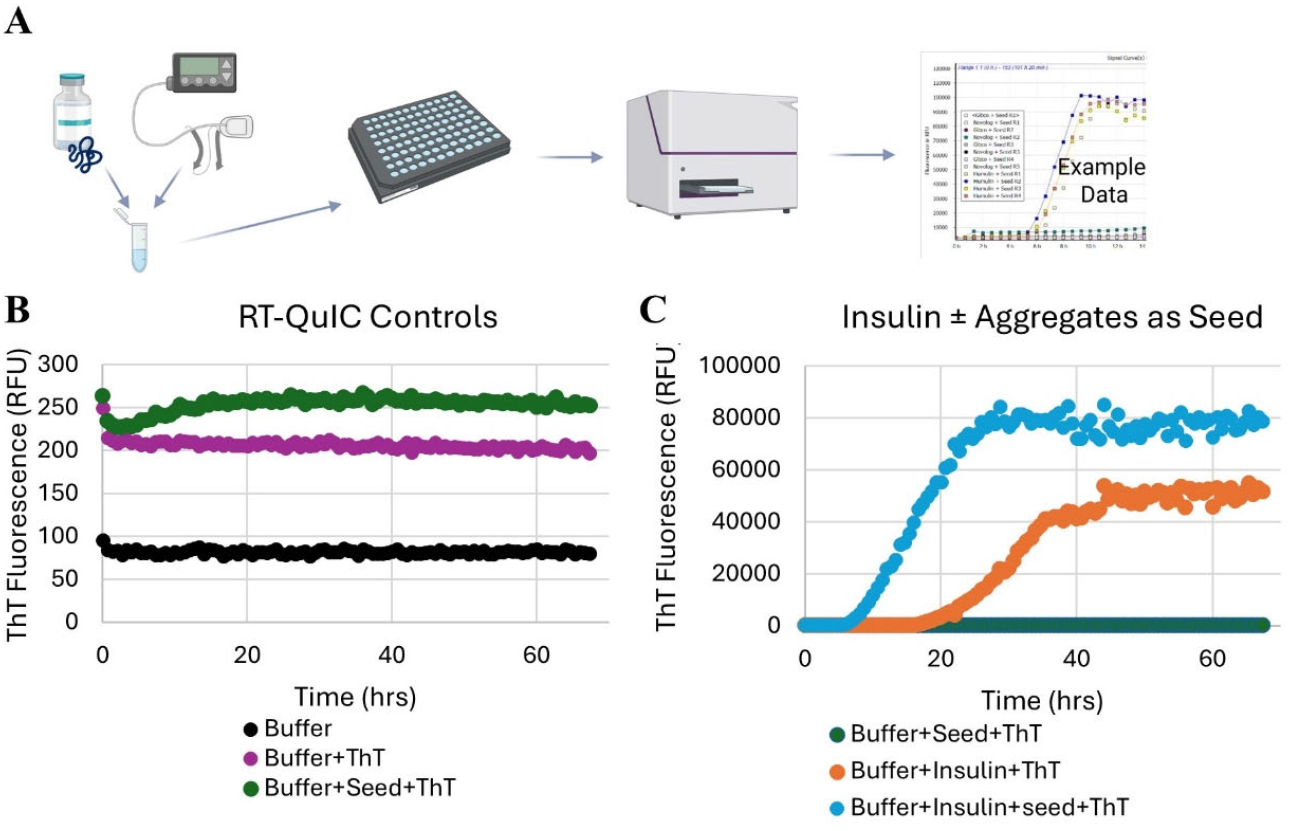
Recombinant Insulin RT-QuIC Assay. **A)** Schematic of the RT-QuIC workflow. Samples arc prepared in master mixes and technical replicates are loaded into 96 well plates then scaled with film. A plate reader incubates the plates at 37°C which undergoes cycles of double orbital shaking followed by rest periods repeatedly for 50-72 boors. ThT fluorescence is measured every 15 minutes. **B)** Representative single sample traces showing RT-QuIC negative controls showing signal levels for the buffer, buffer and fluorescence marker (ThT) and buffer with insulin with no ThT. **C)** Single trace samples of *in vitro* insulin RT-QuIC assay showing signals for non-aggregated insulin alone, non-aggregated insulin with ThT. and non-aggregated insulin with seed (aggregated insulin) and ThT. Insulin alone (blue) has a delayed aggregation profile relative to insulin with seed added (orange).

RT-QuIC was also performed with BSA and lysozyme proteins with no seed, which both had no increase in ThT signal above negative controls (**Supplemental Figure 4**). We also investigated whether BSA or lysozyme could be used as seed to induce the aggregation of recombinant insulin. Neither BSA or lysozyme seeds increased the RT-QuIC aggregation kinetics of recombinant insulin compared to insulin alone, all displaying similar time to half maximum fluorescence signal. **(Supplemental Figure 5)**.

**Figure 5:**
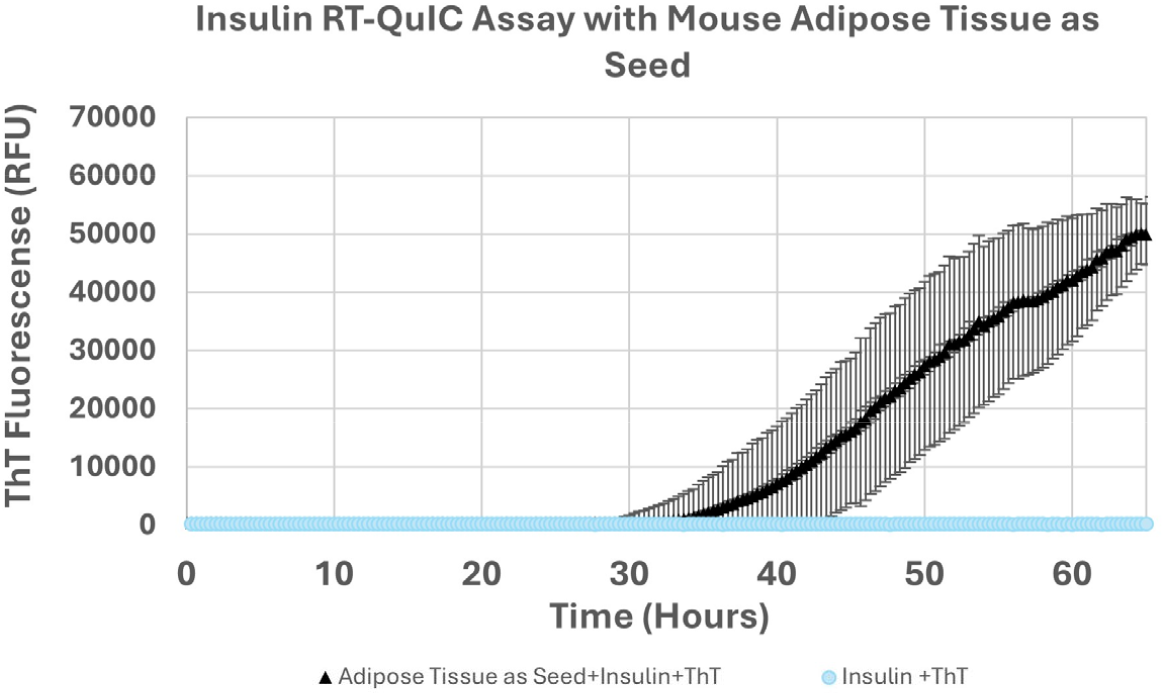
Insulin RT-QuIC in post-mortens mouse tissue. RT-QuIC samples with homogenized adipose (1:10^−6^) tissue from 6-month old J:ARC (S) mice added as seed to recombinant insulin and ThT.

### 3.4 Detection of Misfolded Insulin in Post-Mortem Mouse Tissue

We furthered our proof-of-concept of the RT-QuIC assay to assess whether peripheral tissues contain misfolded protein that could promote aggregation of the recombinant insulin substrate. We collected adipose tissue from J: ARC(S) e, an outbred mouse line that develops progressive weight gain, a primary risk factor for insulin dysregulation. Homogenized mouse adipose tissue was diluted in 1X PBS, pH 7.4 and added as the seed to recombinant, non-aggregated insulin serving as the substrate. Adipose mouse tissue as seed produced aggregated insulin substrate when undergoing the RT-QuIC assay **(Figure 5**). The adipose mouse tissue generated a strong fluorescence signal beginning at 30 hours. Additionally, a total of three mouse adipose tissue dilutions were tested as seed in the RT-QuIC assay and promoted the aggregation of the recombinant insulin in a dilution specific manner. (**Supplemental Figure 6)**.

## 4. Discussion

In this study, we established the feasibility of using RT-QuIC to detect misfolded aggregated insulin, offering a potential new tool for investigating insulin misfolding in the context of metabolic disease. RT-QuIC assays rely on the amplification of a misfolded protein signal based on the principle that native substrate protein will be converted into amyloidized structures in the presence of seeds faster than they would spontaneously nucleate into beta sheet formations without seed. The misfolded protein nucleation seeds amplify the signal of misfolded protein by recruiting normal recombinant protein. To date, sensitive seed amplification assays, such as RT-QuIC, have been used to detect pathogenic proteins in neurodegenerative diseases [24,43,44]. Researchers most often study misfolded proteins in the context of neurodegenerative disorders such as Alzheimer’s, Parkinson’s, and prion diseases, where evidence strongly suggests that these proteins play pathological roles, even beyond the nervous system [41,45,46]. Misfolding and aggregation disrupt fundamental cellular processes, with evidence increasingly indicating a contribution to systemic diseases [47-49]. Despite this evidence, the role of protein misfolding in non-neurodegenerative conditions is less widely studied, particularly in metabolic diseases such as diabetes mellitus. The work presented here is the first to establish suitable seed material and demonstrate the application of RT-QuIC for detecting insulin misfolding.

RT-QuIC assays overcome the limitations of traditional protein detection methods which rely heavily on antibody-based methods, such as Western Blotting, ELISA (enzyme-linked immunosorbent assays), or immunohistochemistry, which utilize specific epitopes of the protein to allow antibody binding. While antibodies can detect prion or other amyloid susceptible protein if the epitope is present, the protein structural configurations are not always distinguishable by antibody -epitope binding. In these cases, researchers may use methods such as proteinase K digestion to degrade the normal non-amyloidized protein, enriching for the more resistant misfolded form. However, the low abundance of aggregated proteins can make detection challenging, and digestion further reduces the amount of target protein. Additionally, mass spectrometry, an alternative method of protein identification, falls short in detecting small amounts of amyloidized proteins because the molecular weights of normal and misfolded proteins are identical. Thus, RT-QuIC can address these challenges by detecting conformational differences rather than relying on sequence or size. RT-QuiC assays permit detection of these misfolded proteins in low concentrations, even leading to antemortem diagnostic tools in the case of prion diseases.

Emerging evidence suggests that insulin misfolding and aggregation may compromise its biological activity, initiating cellular stress responses that contribute to metabolic dysfunction [50,51]. In adding to the limited literature base for this novel view, we developed and validated a prototype screening assay to detect misfolded insulin. The RT-QuIC assay amplifies trace amounts of misfolded ʺseedʺ protein through repeated cycles of shaking and incubation, triggering a conformational change in recombinant insulin substrate. Our study utilized recombinant insulin as the substrate and aggregated insulin recovered from clogged insulin pumps as the seed. We monitored this structural shift in real time using ThT, a fluorescent dye that binds specifically to β-sheet-rich aggregates, enabling quantitative analysis of misfolding kinetics both *in vitro* and in tissues.

In future studies, we aim to apply this assay to cultured cells and animal models exhibiting insulin resistance and chronic hyperglycemia. Our goal is to determine whether insulin misfolding occurs in these systems, whether it correlates with disease severity, and which molecular pathways drive the misfolding process. These studies hold promise of further clarifying whether misfolded insulin acts as a causative factor, a disease biomarker, or a consequence of diabetes-related cellular stress. Through this work, we can extend the detection of misfolded proteins beyond neurodegenerative contexts and into the study of metabolic diseases. If successful, our assay could provide new strategies for early detection, mechanistic insights, and therapeutic interventions in diabetes and related disorders.

## 5. Conclusions

This study demonstrates the feasibility of adapting RT-QuIC as a sensitive and specific approach for detecting misfolded, aggregated insulin, expanding the application of seed amplification assays beyond their traditional use in neurodegenerative diseases. By leveraging confirmational amplification rather than epitope recognition or molecular weight, RT-QuIC overcomes key limitations of conventional protein detection methods and enables the identification of low-abundance misfolded insulin species. Our establishment of suitable seed materials and recombinant-insulin based assay provides a foundational framework for studying insulin misfolding.

These findings support a growing paradigm in which protein misfolding contributes to systemic disease including metabolic disorders such as diabetes. The development of this assay opens new avenues to investigate whether insulin misfolding plays a causal, correlative, or consequential role in disease pathogenesis. Future applications in cellular and *in vivo* models will be critical to defining the biological and clinical significance of these aggregates. Ultimately this work lays the groundwork for extending misfolded protein detection into metabolic disease research with potential implications for early diagnosis, mechanistic understanding and therapeutic innovation.

## 6. Patents

This work is disclosed in PCT Application No. PCT/US2025/053479 filed on October 31, 2025.

## Supplementary Materials

The following supporting information can be downloaded at: https://www.mdpi.com/article/doi/, Figure S1: SDS-PAGE of recombinant non-aggregated insulin and aggregated insulin from an insulin pump; Figure S2: Characterization of Insulin RT-QuIC assay with and without aggregated insulin as seed and recombinant insulin as substrate; Figure S3: Characterization of insulin aggregation kinetics in RT-QuIC by Time to Half Maximum Fluorescence; Figure S4: RT-QuIC Insulin aggregation compared to Bovine Serum Albumin and Lysozyme Protein Aggregation; Figure S5: Insulin RT-QuIC using Alternative Proteins as Seed Materials to Investigate Insulin Aggregation; Figure S6: Dilution series of mouse adipose tissue homogenate as seed for insulin RT-QuIC assay.

## Author Contributions

Jessica M Alderiso: Data curation, Investigation, Methodology, Writing – original draft, Writing – review & editing. Keileigh Fulbright: Data curation, Investigation, Project administration, Supervision, Writing – review & editing. Matthew G DiGiovanni: Conceptualization, Methodology, Writing – review & editing. Rafael Hernandez Latorre: Investigation, Methodology, Validation, Writing – review & editing. Trisha M Cox: Data curation, Investigation, Writing – review & editing. Brenda F Canine: Conceptualization, Formal Analysis, Funding acquisition, Investigation, Methodology, Project administration, Resources, Software, Supervision, Validation, Visualization, Writing – original draft, Writing – review & editing. All authors have read and agreed to the published version of the manuscript.

## Funding

Research was funded by National Institute of General Medical Sciences of the National Institutes of Health under Award Number P20GM152335 and by the Office of Research Infrastructure Programs of the National Institutes of Health under award number 1S10OD038298-01. Jessica Alderiso and Rafael Hernandez were supported by the Touro COM-MT federal work study program

## Institutional Review Board Statement

The research presented here was approved by the McLaughlin Research Institute; McLaughlin Research Institutional Animal Care and Use Committee under protocol number 2025-RP-

## Data Availability Statement

Raw datasets generated during the current study are available from the corresponding author upon reasonable request.

## Acknowledgments

We would like to thank the COBRE administrative core for discussions regarding this manuscript. Colony management for the mouse models included in this study was provided by the McLaughlin Research Institute -Gene Editing and Mouse Models Assessment (GEMMA) Core Facility within the Center for Integrated Biomedical and Rural Health Research, RRID:SCR_027045, 1P20GM152335.

## Conflicts of Interest

The authors declare no conflicts of interest.

## Abbreviations

The following abbreviations are used in this manuscript:

RT-QuIC: Real-time quaking induced conversion
T2DM: Type 2 diabetes mellitus
ATP: Adenosine triphosphate
GDP: Guanosine diphosphate
ThT: Thioflavin T
ALS: Amyotrophic lateral sclerosis
IR: Insulin receptor
IRS1: Insulin receptor substrate 1
IRS2: Insulin receptor substrate 2
PI3K: Phosphoinositide 3-kinase
PIP2: Phosphatidylinositol 3,4,5,-triphosphate
PDK: 3-phsphoinositidtide -dependent kinase-1
Akt: Protein kinase B
GLUT4: Glucose transporter 4
mTOR: Mechanistic target of rapamycin
SDS-PAGE: Sodium dodecyl sulfate-polyacrylamide gel electrophoresis
DPBS: Dulbecco’s phosphate buffered saline
UV: Ultraviolet
BeStSel: Beta Structure Selection methods
CD: Circular dichroism
DLS: Dynamic light scattering
NaCl: Sodium chloride
BSA: Boine serum albumin
RPM: Revolutions per minute
RFUs: Relative fluorescence units
PBS: Phosphate buffered saline
ELISA: Enzyme-linked immunosorbent assay
GEMMA: Gene-Editing and Mouse Models Assessment Core Facility

## Disclaimer/Publisher’s Note

The statements, opinions and data contained in all publications are solely those of the individual author(s) and contributor(s) and not of MDPI and/or the editor(s). MDPI and/or the editor(s) disclaim responsibility for any injury to people or property resulting from any ideas, methods, instructions or products referred to in the content.

---

**Supplemental Figure 1:**
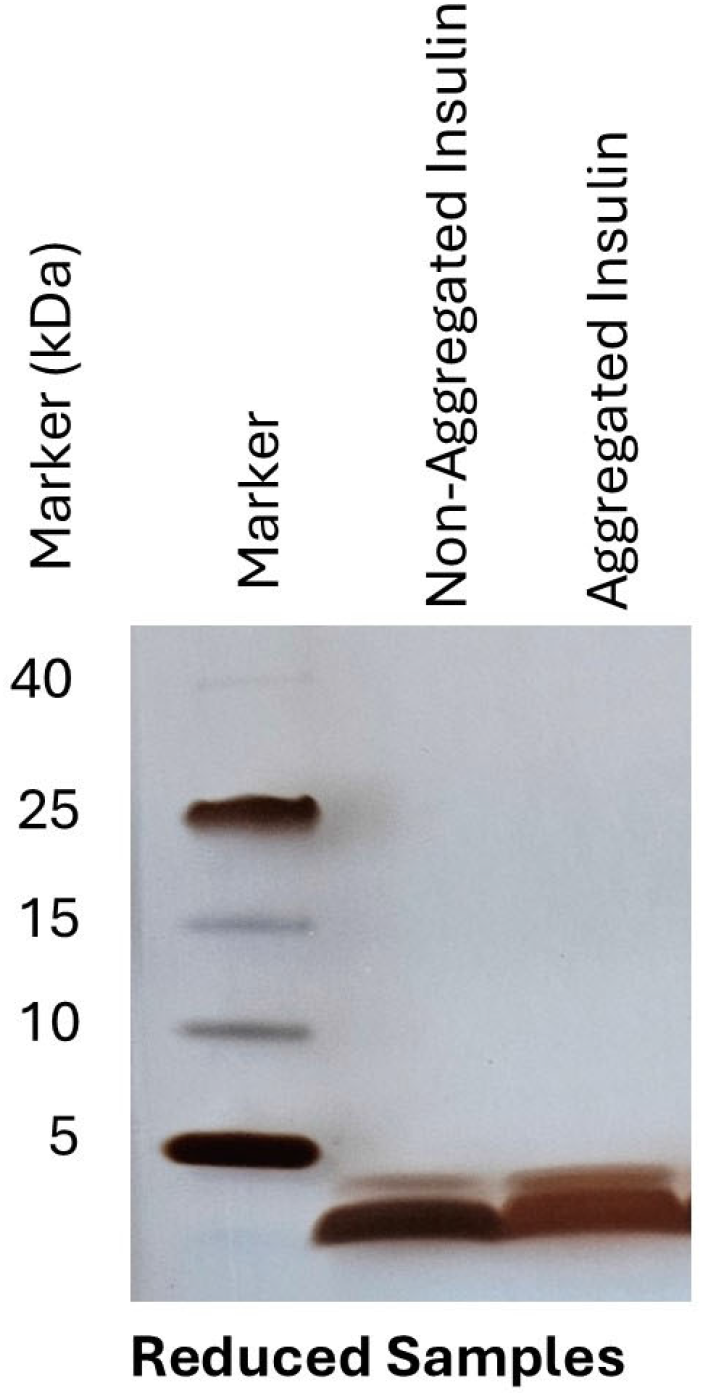
SDS-PAGE of recombinant non-aggregated insulin and aggregated insulin from an insulin pump. A 10-20% Tricine SDS-PAGE was run under reduced conditions to resolve insulin. Protein bands were visualized with silver stain. Molecular weight markers are indicated and ranged from 40 kDa to 5.0 kDa.

**Supplemental Figure 2:**
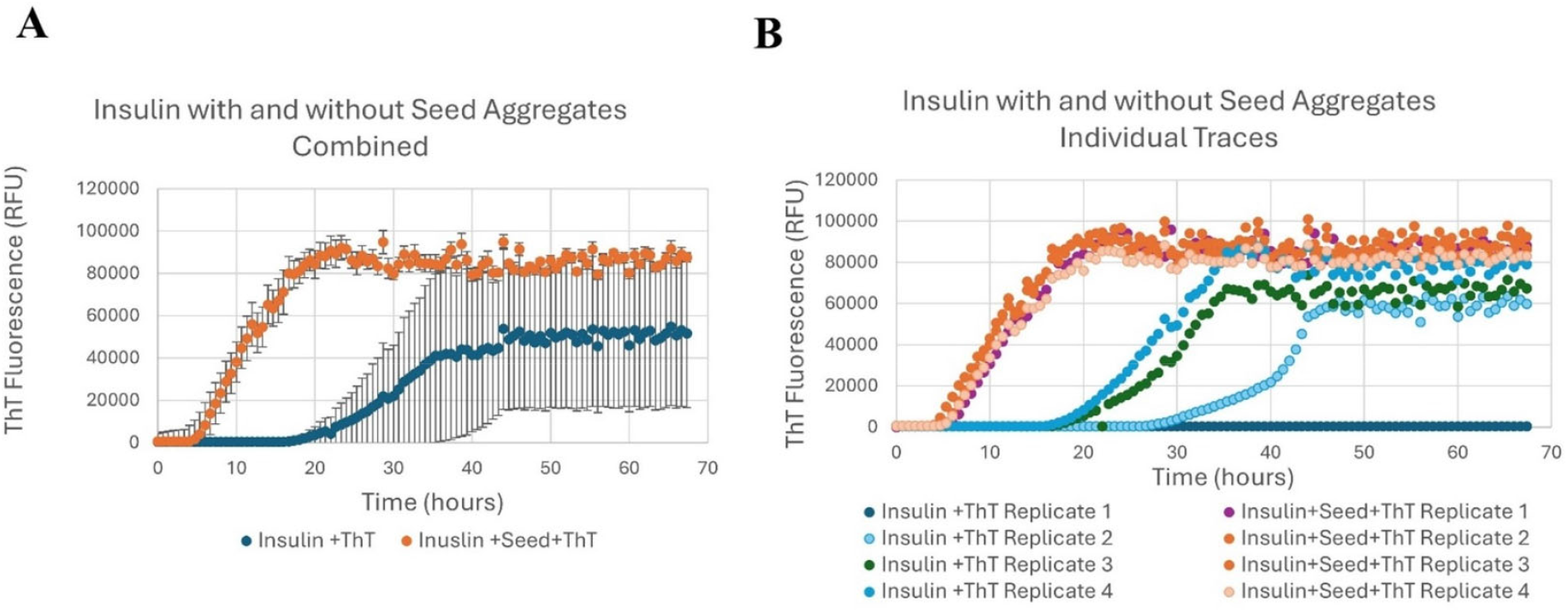
Characterization of Insulin RT-QuIC assay with and without aggregated insulin as seed and recombinant insulin as substrate. **A)** Mean ThT fluorescence ± SD (n=4) of non-aggregated insulin as substrate into the RT-QuIC assay with and without aggregated insulin as seed. **B)** Individual traces of replicates plotted in A.

**Supplemental Figure 3:**
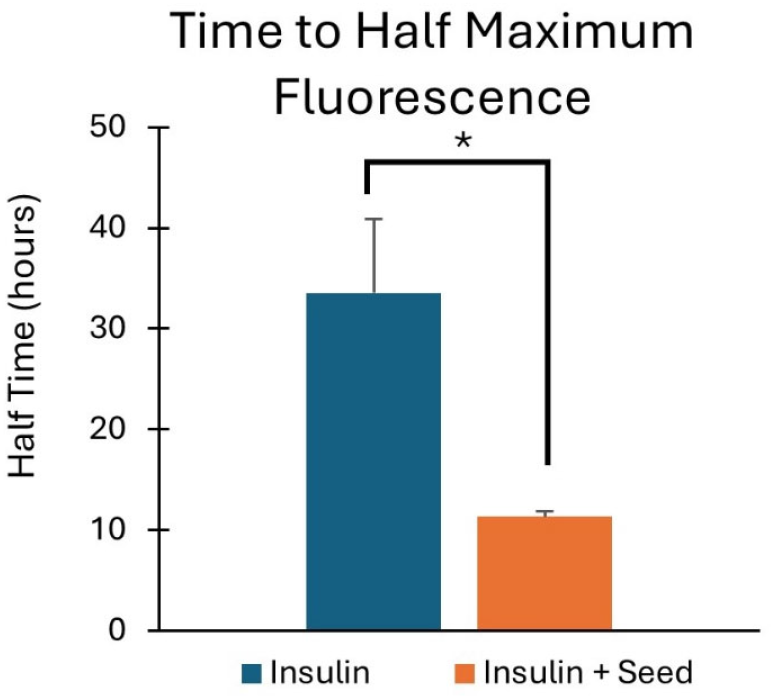
Characterization of insulin aggregation kinetics in RT-QuIC by Time to Half Maximum Fluorescense. For each condition shown in Supplemental Figure 2B the average time to half the maximum ThT signal is displayed± SEM. Significant differences were observed in the average half time of insulin alone compared to insulin with seed aggregates when assayed using RT-QuIC (p = 0.000189).

**Supplemental Figure 4:**
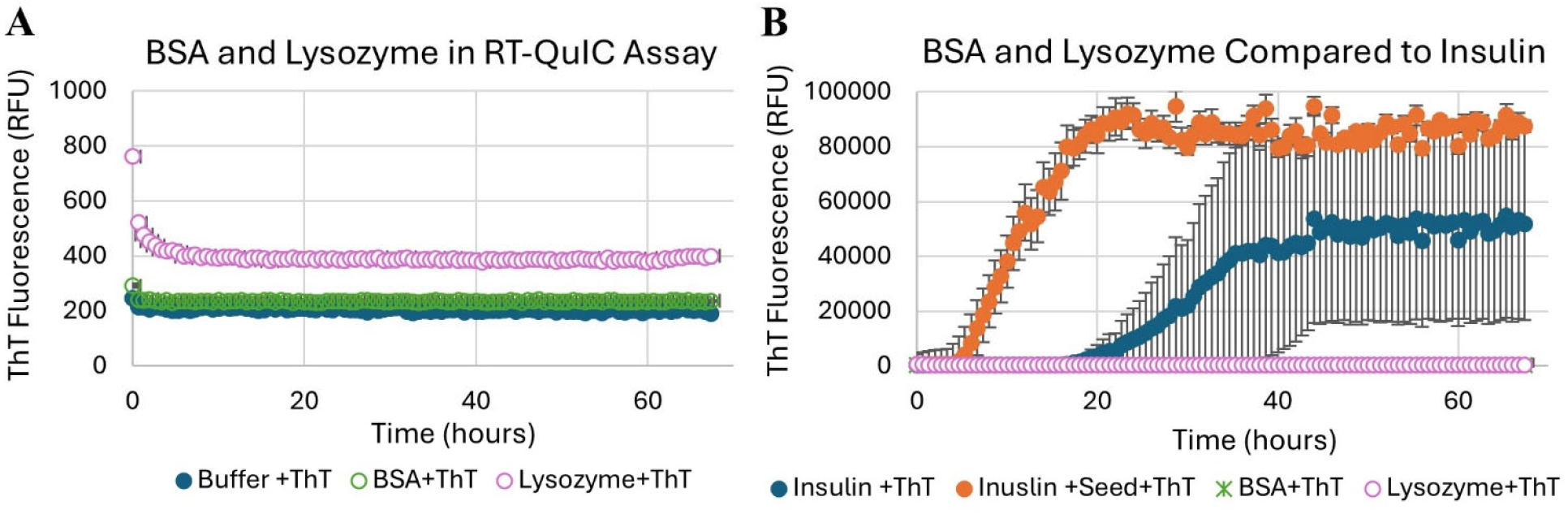
RT-QuIC Insulin aggregation compared to Bovine Serum Albumin (BSA) and Lysozyme Protein Aggregation. **A)** Mean ThT Fluorescence± SD (n=4) of BSA and Lysozyme as substrate for the RT-QuIC assay. **B)** Mean ThT Fluorescence ± SD (n=4) comparison between RT-QuIC substrates to assess aggregation: BSA, Lysozyme, insulin, and insulin with aggregated insulin as seed.

**Supplemental Figure 5:**
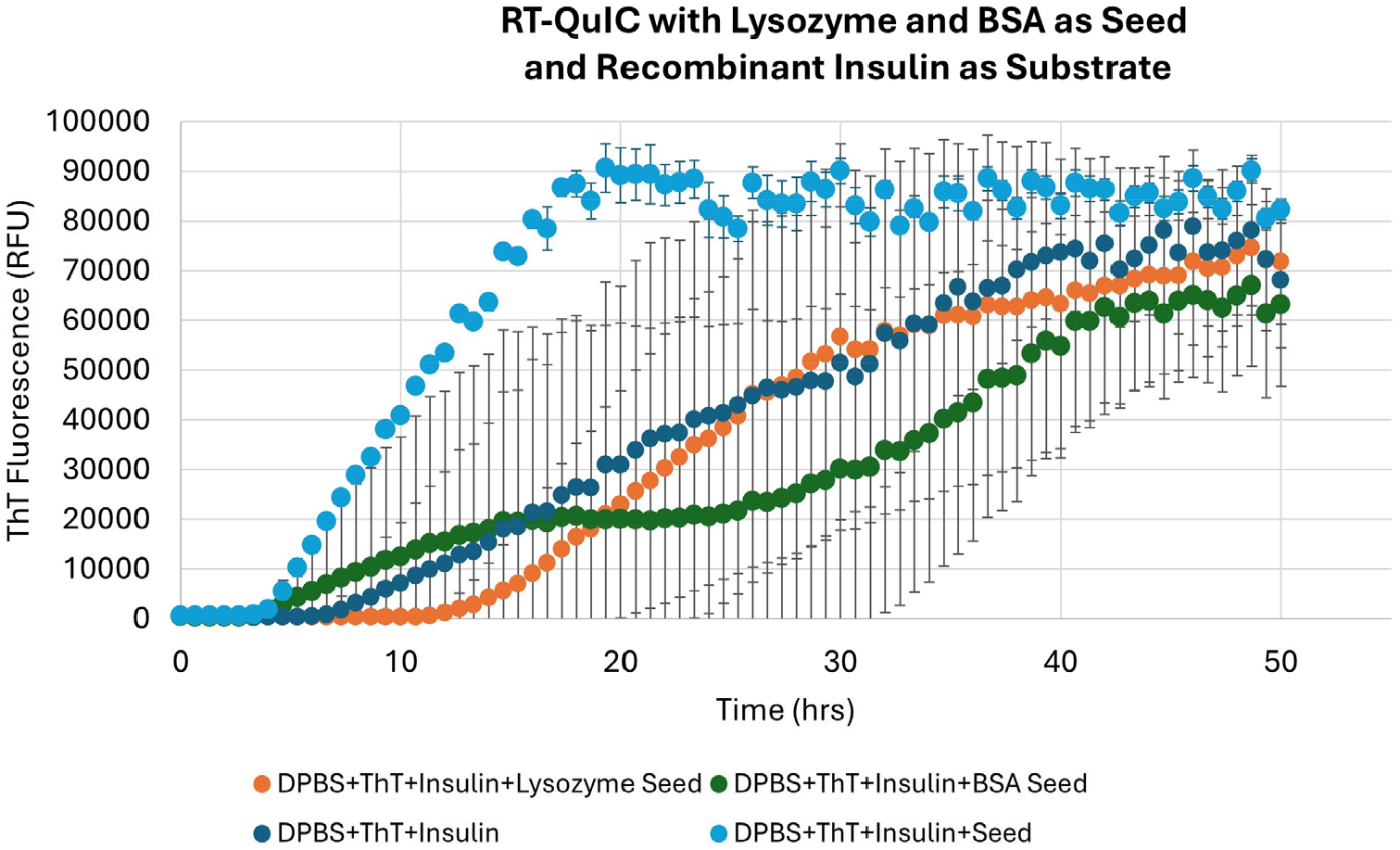
Insulin RT-QuIC assay using Alternative Proteins as Seed Materials to Investigate Insulin Aggregation. RT-QuIC assay was performed as described previously with recombinant non-aggregated insulin as the substrate material. Instead of aggregated insulin being added as a seed material, BSA or Lysosome in equivalent amounts was added to the recombinant non-aggregated insulin. Mean ThT Fluorescence± SD (n=4) are displayed.

**Supplemental Figure 6:**
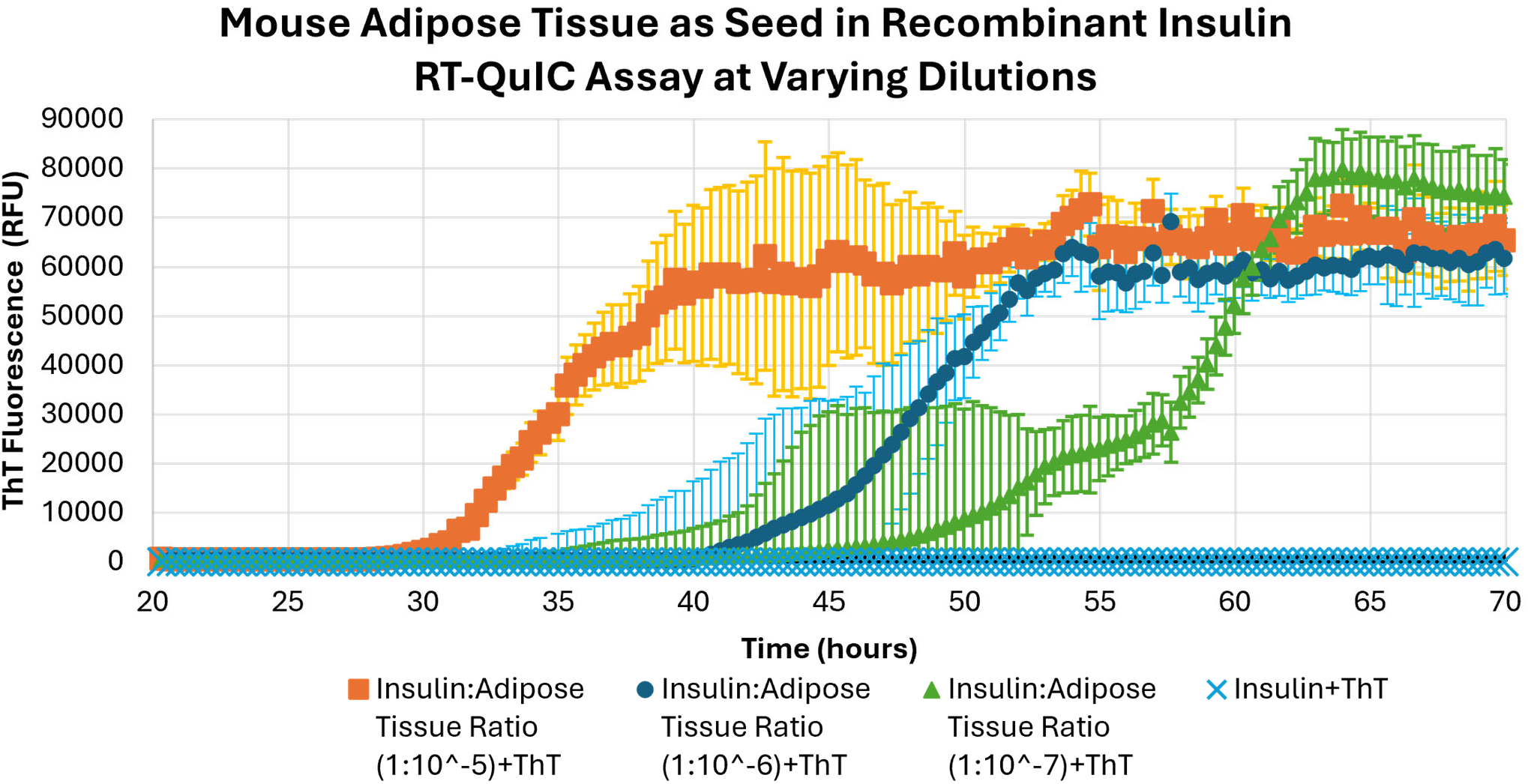
Dilution series of mouse adipose tissue homogenate as seed for insulin RT-QuIC assay. Homogenized mouse adipose was diluted to 1:10^− 5^, 1:10^−6^, and l:10^−7^in lX PBS (pH 7.4) and input as seed material for RT-QuIC assays using non-aggregated insulin as substrate. Mean ThT Fluorescence± SD (n=4) are displayed.

## Notes

### Competing Interest Statement

The authors have declared no competing interest.

